# Genetic Associations with Subjective Well-Being Also Implicate Depression and Neuroticism

**DOI:** 10.1101/032789

**Authors:** Aysu Okbay, Bart M.L. Baselmans, Jan-Emmanuel De Neve, Patrick Turley, Michel G. Nivard, Mark A. Fontana, S. Fleur W. Meddens, Richard Karlsson Linnér, Cornelius A. Rietveld, Jaime Derringer, Jacob Gratten, James J. Lee, Jimmy Z. Liu, Ronald de Vlaming, Tarunveer S. Ahluwalia, Jadwiga Buchwald, Alana Cavadino, Alexis C. Frazier-Wood, Nicholas A. Furlotte, Victoria Garfield, Marie Henrike Geisel, Juan R. Gonzalez, Saskia Haitjema, Robert Karlsson, Sander W. van der Laan, Karl-Heinz Ladwig, Jari Lahti, Sven J. van der Lee, Penelope A. Lind, Tian Liu, Lindsay Matteson, Evelin Mihailov, Michael B. Miller, Camelia C. Minica, Ilja M. Nolte, Dennis Mook-Kanamori, Peter J. van der Most, Christopher Oldmeadow, Yong Qian, Olli Raitakari, Rajesh Rawal, Anu Realo, Rico Rueedi, Börge Schmidt, Albert V. Smith, Evie Stergiakouli, Toshiko Tanaka, Kent Taylor, Juho Wedenoja, Juergen Wellmann, Harm-Jan Westra, Sara M. Willems, Wei Zhao, Behrooz Z. Alizadeh, Najaf Amin, Andrew Bakshi, Patricia A. Boyle, Simon Cox, Gail Davies, Oliver S.P. Davis, Jun Ding, Nese Direk, Peter Eibich, Rebecca T. Emeny, Ghazaleh Fatemifar, Jessica D. Faul, Luigi Ferrucci, Andreas J. Forstner, Christian Gieger, Tamara B. Harris, Juliette M. Harris, Elizabeth G. Holliday, Jouke-Jan Hottenga, Philip L. De Jager, Marika A. Kaakinen, Eero Kajantie, Ville Karhunen, Ivana Kolcic, Meena Kumari, Lenore J. Launer, LifeLines Cohort Study, Ruifang Li-Gao, Marisa Loitfelder, Anu Loukola, Pedro Marques-Vidal, Grant W. Montgomery, Lavinia Paternoster, Alison Pattie, Katja E. Petrovic, Laura Pulkki-Råback, Lydia Quaye, Katri Räikkönen, Igor Rudan, Rodney J. Scott, Jennifer A. Smith, Angelina R. Sutin, Maciej Trzaskowski, Anna E. Vinkhuyzen, Lei Yu, Delilah Zabaneh, John R. Attia, David A. Bennett, Klaus Berger, Lars Bertram, Dorret I. Boomsma, Ute Bultmann, Shun-Chiao Chang, Francesco Cucca, Ian J. Deary, Cornelia M. van Duijn, Johan G. Eriksson, Lude Franke, Eco J.C. de Geus, Patrick J.F. Groenen, Vilmundur Gudnason, Torben Hansen, Catharine A. Hartman, Claire M.A. Haworth, Caroline Hayward, Andrew C. Heath, David A. Hinds, Elina Hyppönen, William G. Iacono, Marjo-Riitta Järvelin, Karl-Heinz Jöckel, Jaakko Kaprio, Sharon L.R. Kardia, Liisa Keltikangas-Järvinen, Peter Kraft, Laura Kubzansky, Terho Lehtimäki, Patrik K.E. Magnusson, Nicholas G. Martin, Matt McGue, Andres Metspalu, Melinda Mills, Renée de Mutsert, Albertine J. Oldehinkel, Gerard Pasterkamp, Nancy L. Pedersen, Robert Plomin, Ozren Polasek, Christine Power, Stephen S. Rich, Frits R. Rosendaal, Hester M. den Ruijter, David Schlessinger, Helena Schmidt, Rauli Svento, Reinhold Schmidt, Harold Snieder, Thorkild I.A. Sørensen, Tim D. Spector, Andrew Steptoe, Antonio Terracciano, A. Roy Thurik, Nicholas J. Timpson, Henning Tiemeier, André G. Uitterlinden, Peter Vollenweider, Gert Wagner, David R. Weir, Jian Yang, Dalton C. Conley, George Davey Smith, Albert Hofman, Magnus Johannesson, David I. Laibson, Sarah E. Medland, Michelle N. Meyer, Joseph K. Pickrell, Tõnu Esko, Robert F. Krueger, Jonathan P. Beauchamp, Philipp D. Koellinger, Daniel J. Benjamin, Meike Bartels, David Cesarini

## Abstract

We conducted a genome-wide association study of subjective well-being (SWB) in 298,420 individuals. We also performed auxiliary analyses of depressive symptoms (“DS”; *N* = 161,460) and neuroticism (*N* = 170,910), both of which have a substantial genetic correlation with SWB 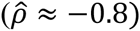. We identify three SNPs associated with SWB at genome-wide significance. Two of them are significantly associated with DS in an independent sample. In our auxiliary analyses, we identify 13 additional genome-wide-significant associations: two with DS and eleven with neuroticism, including two inversion polymorphisms. Across our phenotypes, loci regulating expression in central nervous system and adrenal/pancreas tissues are enriched. The discovery of genetic loci associated with the three phenotypes we study has proven elusive; our findings illustrate the payoffs from studying them jointly.

**One Sentence Summary**: Using both genome-wide association studies and proxy-phenotype studies, we identify genetic variants associated with subjective well-being, depressive symptoms, and neuroticism.

---

**Main Text**: Subjective well-being (SWB)—as measured by survey questions on life satisfaction, positive affect, or happiness—is a major topic of research within psychology, economics, and epidemiology and is the focus of policy initiatives instigated by many governments and international bodies (*1–3*). Twin studies suggest that roughly 35% of the variation in SWB across individuals can be explained by genetic factors (*4*). The discovery of genetic variants associated with SWB could pave the way for investigations into how environmental conditions moderate genetic effects. Moreover, it may provide insights not only into the molecular mechanisms underlying normal-range variation in SWB, but also those underlying genetically correlated phenotypes, such as depression and neuroticism (*5–7*), for which the discovery of genetic associations has proven elusive (*8, 9*).

Here, we report a series of analyses that exploit the relationships between SWB, depressive symptoms (DS), and neuroticism. Our primary analysis is a genome-wide association study (GWAS) of SWB based on data from 59 cohorts (*N* = 298,420). We supplement this primary analysis with auxiliary GWAS meta-analyses of DS and neuroticism. The auxiliary analyses are performed by combining publicly available summary statistics from published studies with new genome-wide analyses of additional data. All analyses in this paper are restricted to European-ancestry individuals.

Following a pre-specified analysis plan, our primary analysis combines survey measures of life satisfaction (LS) and positive affect (PA). Although LS and PA are distinguishable (*10, 11*), these two facets of SWB are phenotypically correlated and load on a common genetic factor (*12*). To reduce measurement error, cohorts with repeated measures averaged responses, and cohorts with survey questions measuring both LS and PA combined the two measures into a single variable. For details on SWB phenotype construction, see tables S1.1 and S2.1.

All SWB analyses were performed at the cohort level and subsequently meta-analyzed. Our analysis plan specified controls for age, age^2^, sex, and (to reduce confounding from population stratification) four principal components of the genome-wide data (*13*). A uniform set of quality-control (QC) procedures, described in tables S2.3-2.4, were applied to the cohort-level summary statistics, including but not limited to the EasyQC protocol recently recommended (*14*).

We conducted a sample-size-weighted meta-analysis of ~2.3M HapMap2 SNPs and adjusted the standard errors using the estimated intercept from an LD Score regression (*15*). The estimated intercept implies that a modest ~5% of the observed inflation in the unadjusted mean *χ^2^* is accounted for by bias rather than polygenic signal (fig. S3.1). The slope estimate implies that our phenotype has a SNP-based heritability estimate of 4.0% (SE = 0.2%; cf. (*16*)).

In our primary analysis of SWB, we found three approximately independent genome-wide significant SNPs (hereafter “lead SNPs”). Fig. 1a shows the Manhattan plot from the main analysis. All three lead SNPs have estimated effects in the range 0.015 to 0.018 standard deviations (SDs) per allele (*R^2^* ≈ 0.01%). For comparison, we calculated, using publicly available survey data from the U.S. Behavioral Risk Factor Surveillance System, that the average difference in SWB between “no” and “yes” responders to “ever diagnosed with a depressive disorder” is 0.66 SDs (table S1.2). We also conducted separate meta-analyses of LS (*N* = 166,205) and PA (*N* = 180,281), using procedures identical to those described above. Consistent with our theoretical calculations (fig. S4.1), these analyses yielded fewer signals across a range of *p*-value thresholds (table S2.6). We found two lead SNPs in the LS analyses (both distinct from the SWB lead-SNPs) and none in the PA analyses.

**Fig. 1.**
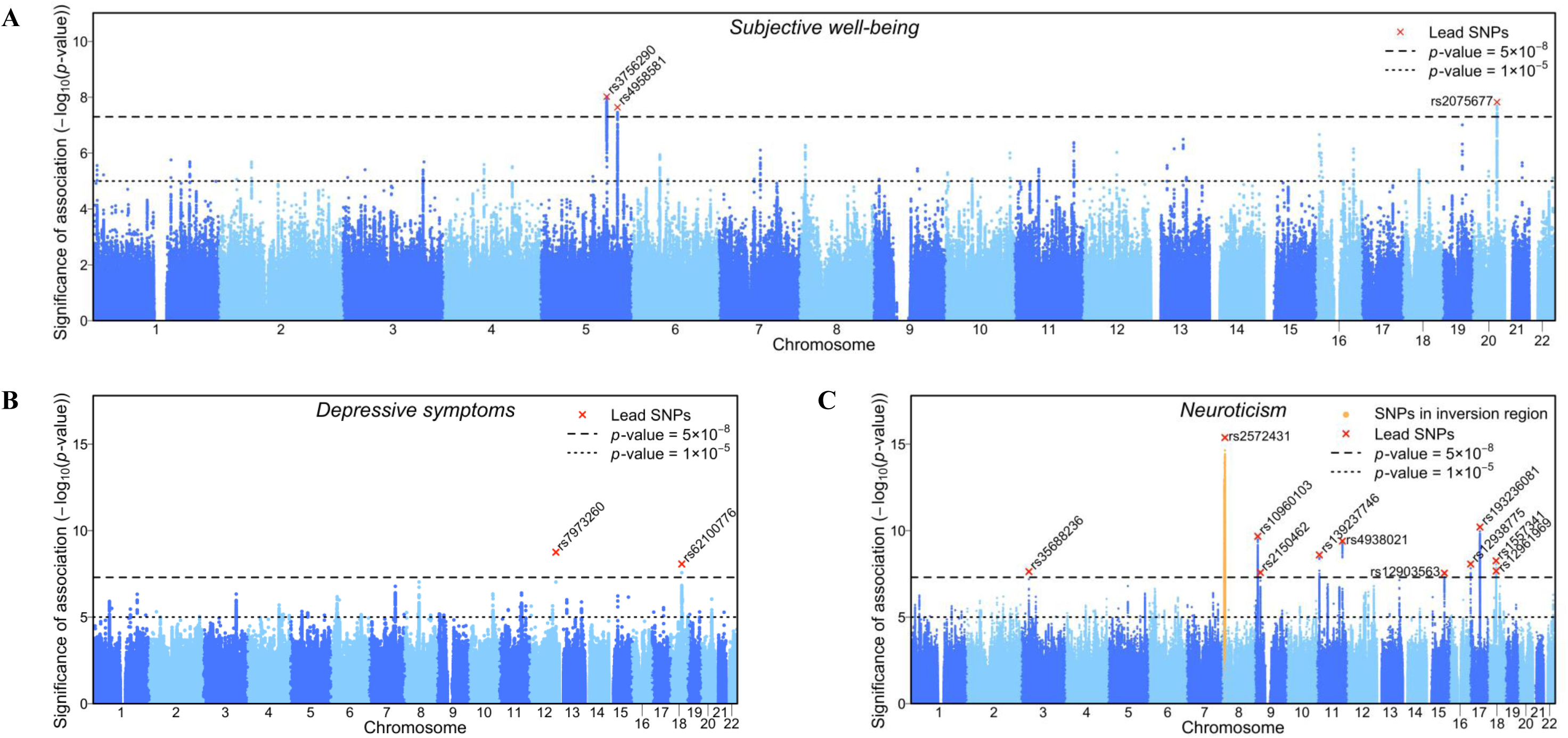
Manhattan plots. **(A)** Subjective well-being (*N* = 298,420), **(B)** Depressive symptoms (*N_eff_* = 149,707), **(C)** Neuroticism (*N* = 170,911). The *x*-axis is chromosomal position, and the *y*-axis is the significance on a −log_10_ scale. The upper dashed line marks the threshold for genome-wide significance (*p* = 5×10^−8^); the lower line marks the threshold for nominal significance (*p* = 10^−5^). Each independent genome-wide significant association (“lead SNP”) is marked by ×. We find three associations for SWB, two for DS, and eleven for neuroticism.

In our auxiliary analysis of DS we combined publicly available results from a study performed by the Psychiatric Genomics Consortium (PGC) (*17*) with new results from analyses of the initial release of the U.K. Biobank data (UKB) and the Resource for Genetic Epidemiology Research on Aging (GERA) Cohort. In UKB (*N* = 105,739), our measure is constructed from responses to two questions about the frequency with which the respondent experienced feelings of hopelessness or depression in the past two weeks. The other cohorts have case-control data on major depressive disorder (GERA: *N_cases_* = 7,231, *N_controls_* = 49,316; PGC: *N_cases_* = 9,240, *N_controls_* = 9,519). In the DS meta-analysis, we weight UKB by sample size and the two case-control studies by effective sample size (*18*).

In our auxiliary analysis of neuroticism (*N* = 170,910), we pooled summary statistics from a study by the Genetics of Personality Consortium (GPC) (*8*) with results from a new analysis of UKB data. In UKB, our measure is the respondent’s score on a 12-item version of the Eysenck Personality Inventory Neuroticism scale (*19*). The GPC harmonized different neuroticism batteries (table S5.2).

Figs. 1b and 1c show the Manhattan plots. For DS, we identified two genome-wide significant SNPs. For neuroticism, our meta-analysis procedure yielded 16 seemingly-independent genome-wide significant SNPs. However, 6 of these SNPs reside within a well-known inversion polymorphism on chromosome 8 (*20*). We established that all genome-wide significant signals in the inversion region are attributable to the inversion (fig. S6.3) and confirmed that the inversion is associated with neuroticism in both the GPC and UKB data. Another SNP, rs193236081, is located on a well-known inversion polymorphism on chromosome 17, and we also established that this association is attributable to the inversion polymorphism (fig S6.6). However, because this inversion yields only one significant SNP and is genetically complex (*21*), we hereafter simply use rs193236081 as its proxy. Our neuroticism GWAS therefore identified 9 lead SNPs and 2 inversion polymorphisms. A concurrent, unpublished neuroticism GWAS using overlapping data reports similar findings (*22*).

The estimated effects of the genome-wide significant associations on DS and neuroticism are similar, with coefficient estimates in the range 0.020 to 0.031 SDs per allele (*R*^2^ ≈ 0.02%). In the UKB cohort we estimated the effect of an additional allele of the chromosome 8 inversion polymorphism on neuroticism to be 0.035 SDs. For the 16 polymorphisms identified across our main and auxiliary analyses—three for SWB, two for DS, and eleven for neuroticism— Bayesian analyses show that for a wide range of priors, the evidence for association is highly credible (fig. S7.1).

In independent samples, a polygenic score constructed from our primary SWB GWAS results explains ~0.9% of the variance in SWB, ~0.5% in DS, and ~0.7% in neuroticism (table S8.1). Applying bivariate LD Score regression (*23*) to our GWAS results, we found substantial pairwise genetic correlations between SWB and our two auxiliary phenotypes: −0.81 (SE = 0.046) for DS (cf., (*24*)) and −0.75 (SE = 0.034) for neuroticism (table S9.1). As a placebo test, we also examined height, which we expected to have much less genetic overlap with SWB (*25*). Indeed, we find that the polygenic score explains ~0.2% of the variance in height, and the genetic correlation with SWB is only 0.07 (SE = 0.028).

Motivated by this evidence of genetic overlap with DS and neuroticism, which is in line with large genetic correlations estimated from twin-family studies (*5, 6*), we conducted a series of “quasi-replication” exercises in which we examined whether (i) the three genome-wide significant SNPs in the SWB analyses are associated with DS and neuroticism, and (ii) the genome-wide significant hits found in the auxiliary GWAS analyses of DS and neuroticism are associated with SWB. To avoid sample overlap, the analyses of the “second-stage” phenotype were always restricted to cohorts that did not contribute to the “first-stage” analysis.

Fig. 2 illustrates the results from these analyses. For interpretational ease, we choose reference alleles so that each SNP’s estimated effect on SWB is negative; that way, its predicted effect on DS and neuroticism is positive. Panel A shows that two out of the three SWB lead-SNPs are significantly associated with DS in the predicted direction (*p* = 0.004 and *p* = 0.001), and the third has the opposite sign and is not significantly associated with DS. For neuroticism, the SWB-increasing allele has the predicted sign for all three SNPs, but none reach significance (Panel B). Panel C shows that the two DS lead-SNPs have the predicted sign for SWB, and one is nominally significant (*p* = 0.04). Finally, of the eleven polymorphisms associated with neuroticism, four have the predicted sign and are significantly associated with SWB, five have the predicted sign and are not significantly associated with SWB, and two have the opposite sign and are not significantly associated with SWB (Panel D). One of the four significant polymorphisms is the SNP tagging the inversion on chromosome 8 (*20*). That SNP’s association with neuroticism (and likely with SWB) is driven by its correlation with the inversion (fig. S6.3).

**Fig. 2.**
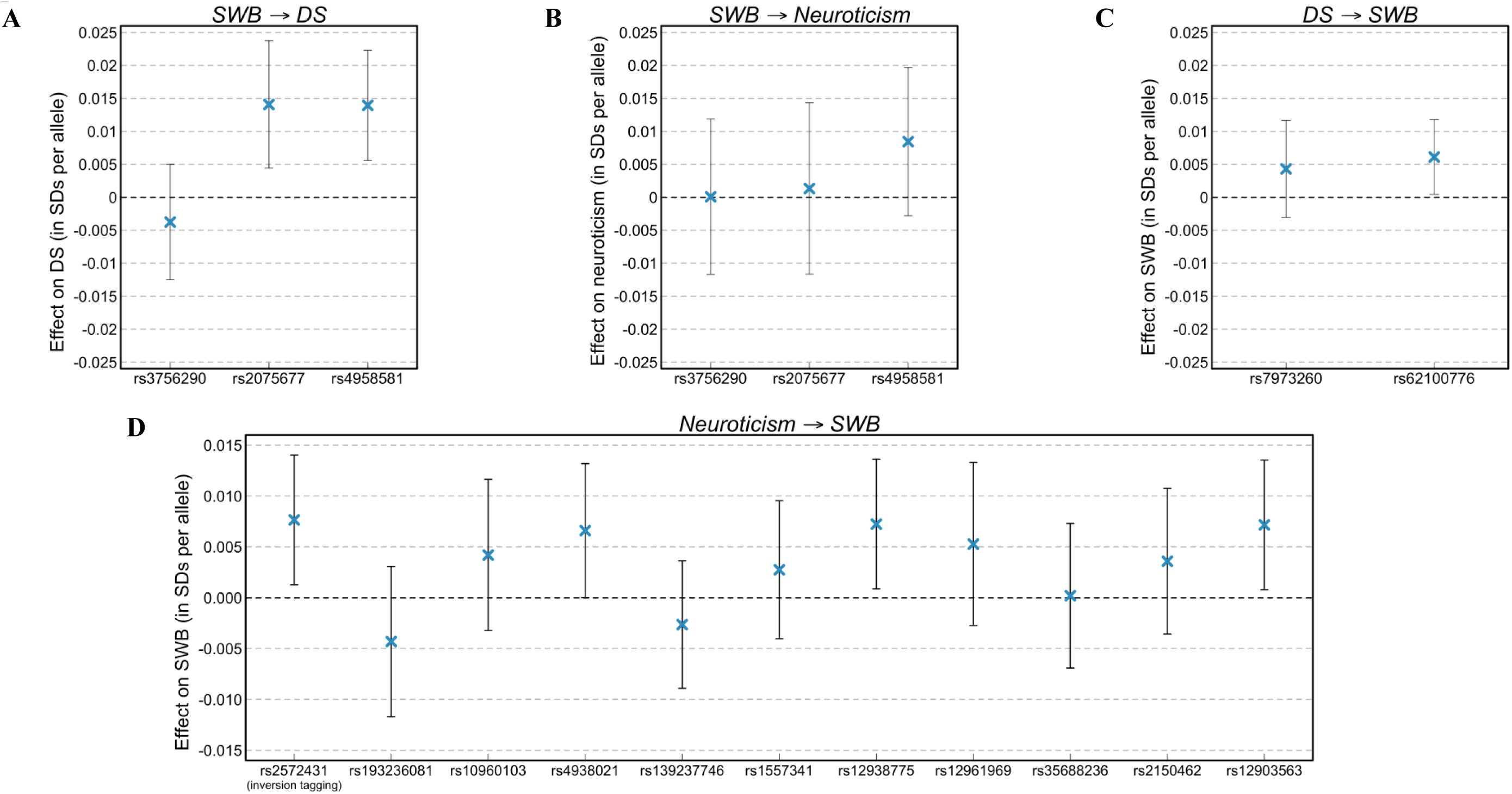
Quasi-replication of lead SNPs. We examined whether (i) lead SNPs identified in the SWB meta-analyses are associated with DS and neuroticism, and (ii) lead SNPs identified in the analyses of DS and neuroticism are associated with SWB. The quasi-replication sample is always restricted to non-overlapping cohorts. The bars represent 95% CIs (not adjusted for multiple testing). The sample sizes (maximum across SNPs) are: **(A)** SWB (*N* = 294,043) → DS (*N_eff_* = 123,506), **(B)** SWB (*N* = 294,043) → Neuroticism (*N* = 68,201), **(C)** DS (*N_eff_* = 124,498) → SWB (*N* = 238,254), **(D)** Neuroticism (*N* = 170,908) → SWB (*N* = 198,360). For interpretational ease, we choose the reference allele so that positive coefficients imply that the estimated effect is in the predicted direction.

Because of the strong genetic correlation between SWB and DS, the two successful quasi-replications of the SWB lead-SNPs for DS lend credibility to their truly being associated with *both* SWB and DS. Indeed, our formal Bayesian calculations (table S11.1) show that for each SNP’s effect on each phenotype, we can exclude a zero effect size with greater than 95% probability. Similarly, across the 19 quasi-replication attempts depicted in fig. 2, with at least 95% probability we can exclude a zero effect size for 18 of the first-stage effects and 9 of the second-stage effects. This Bayesian analysis also yields effect-size estimates that are corrected for the winner’s curse: relative to the initial estimates, on average these estimates are smaller by a factor of four for the first-stage phenotype and two for the second-stage phenotype.

We also conducted two similar “proxy-phenotype” analyses (*26*). We examine whether the independent SNPs associated with SWB at *p* < 10^−4^ and available in the second-stage sample—163 SNPs for DS and 170 for neuroticism—are enriched for association with DS and neuroticism. For both second-stage phenotypes, the observed level of enrichment is stronger than the expected level of enrichment for a randomly drawn set of SNPs matched on allele frequency and second-stage sample size (*p* < 3×10^−5^ and *p* < 0.04, respectively). For DS, 116 out of 163 SNPs (71%) have the predicted direction of effect. Twenty of the 163 SNPs are significantly associated with DS at the 5% level (19 with the predicted direction), and 2 remain significant after Bonferroni correction (fig. S10.1). For neuroticism, 129 out of 170 SNPs (76%) have the predicted direction of effect, all 28 significant SNPs have the predicted sign, and 4 remain significant after Bonferroni correction (fig. S10.2). As a placebo test, we also test SWB-associated SNPs for enriched association with height. In stark contrast to what we find for DS and neuroticism, we find no evidence for enrichment and cannot reject the null hypothesis that the effects have the predicted sign only 50% of the time (*25*).

Table 1 gives a summary overview of the SNPs identified across our analyses. In the upper panel, we list all lead SNPs from the meta-analyses of SWB, DS, or neuroticism, and we indicate which have significant associations in the quasi-replication analyses. In the lower panel, we list the SNPs identified in the proxy-phenotype analyses. For each SNP, we report its estimated effect. Finally, we also report if each SNP (or a variant in strong linkage disequilibrium with that SNP) is (i) nonsynonymous (i.e., alters the protein product) or is (ii) an eQTL (i.e., affects the extent to which certain genes are expressed), and we report (iii) the nearest protein-coding gene; for details, see tables S2.6, S10.2-10.3, S12.5-12.6, and S12.8.

**Table 1.**
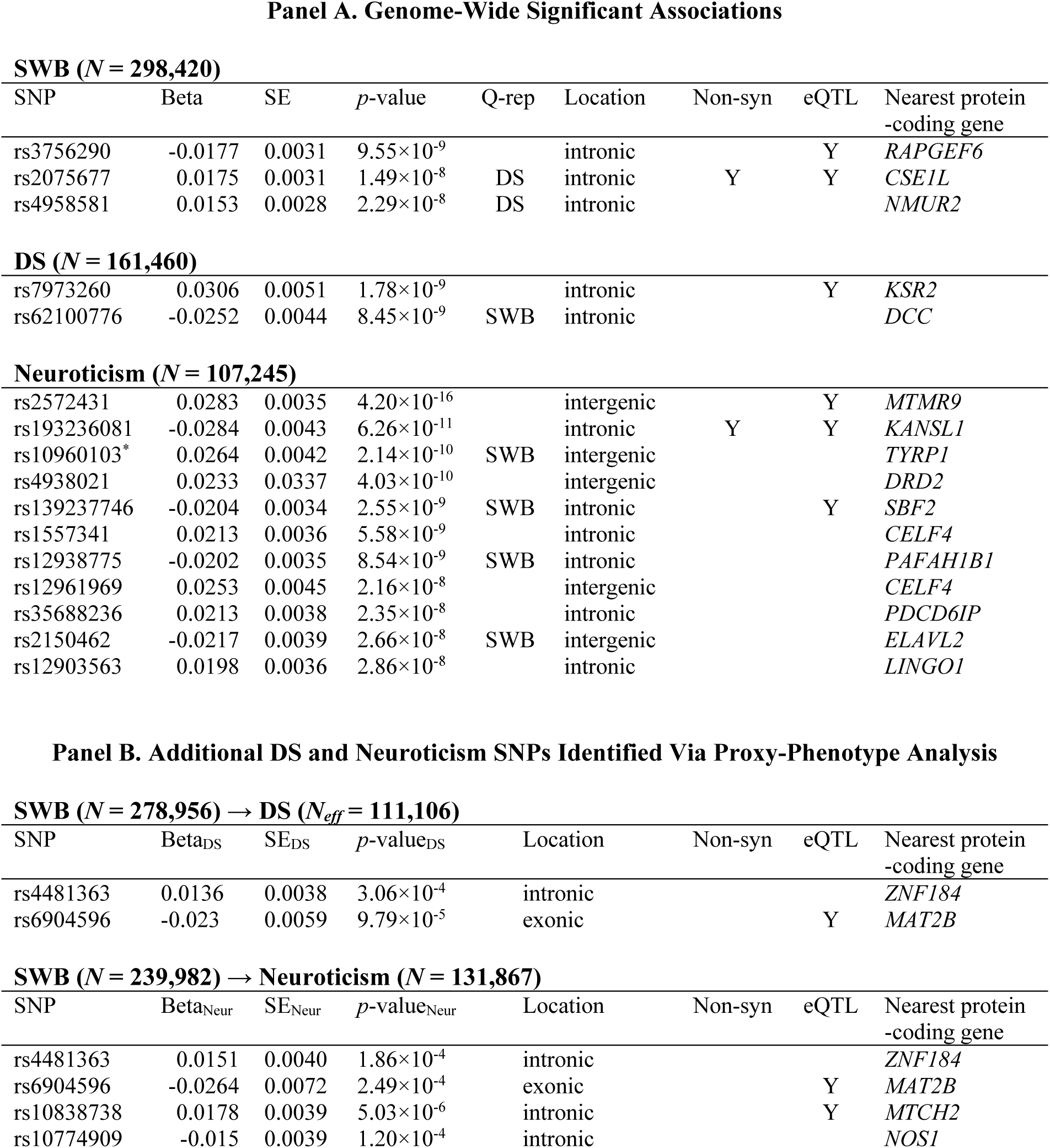
Summary Overview of Polymorphisms Identified Across Analyses. All effect sizes are reported in units of SDs per allele. “Q-rep”: Whether SWB lead-SNP quasi-replicated in DS or neuroticism, and whether DS/neuroticism lead-SNP quasi-replicated in SWB, all at *p* < 0.05. “Location”: type of genomic region according to dbsnp and Ensembl. “Non-syn”: Whether the SNP or any LD partner (*R*^2^ ≥ 0.6 and within 250kb) is non-synonymous according to HaploReg (version 4). “eQTL”: Whether the SNP is a *cis*-eQTL in whole blood (at FDR < 0.05). “Nearest protein-coding gene”: The nearest-protein coding gene (5’ start site) of the lead SNP according to GENCODE coordinates archived at http://www.broadinstitute.org/mpg/snpsnap/. Proxy SNP rs4346787 (*R*^2^ = 0.98) was used to tag rs4481363 in the DS data. All details are in SOM Section 12.2. * Note: rs10960103 tags the inversion polymorphism on chromosome 8.

**SWB ( $N = 298,420$ )**
| SNP | Beta | SE | $p$-value | Q-rep | Location | Non-syn | eQTL | Nearest protein-coding gene |
| --- | --- | --- | --- | --- | --- | --- | --- | --- |
| rs3756290 | -0.0177 | 0.0031 | $9.55 \times 10^{-9}$ | | intronic | | Y | *RAPGEF6* |
| rs2075677 | 0.0175 | 0.0031 | $1.49 \times 10^{-8}$ | DS | intronic | Y | Y | *CSE1L* |
| rs4958581 | 0.0153 | 0.0028 | $2.29 \times 10^{-8}$ | DS | intronic | | | *NMUR2* |
| **DS ( $N = 161,460$ )** |  |  |  |  |  |  |  |  |
| rs7973260 | 0.0306 | 0.0051 | $1.78 \times 10^{-9}$ | | intronic | | Y | *KSR2* |
| rs62100776 | -0.0252 | 0.0044 | $8.45 \times 10^{-9}$ | SWB | intronic | | | *DCC* |
| **Neuroticism ( $N = 107,245$ )** |  |  |  |  |  |  |  |  |
| rs2572431 | 0.0283 | 0.0035 | $4.20 \times 10^{-16}$ | | intergenic | | Y | *MTMR9* |
| rs193236081 | -0.0284 | 0.0043 | $6.26 \times 10^{-11}$ | | intronic | Y | Y | *KANSL1* |
| rs10960103* | 0.0264 | 0.0042 | $2.14 \times 10^{-10}$ | SWB | intergenic | | | *TYRP1* |
| rs4938021 | 0.0233 | 0.0337 | $4.03 \times 10^{-10}$ | | intergenic | | | *DRD2* |
| rs139237746 | -0.0204 | 0.0034 | $2.55 \times 10^{-9}$ | SWB | intronic | | Y | *SBF2* |
| rs1557341 | 0.0213 | 0.0036 | $5.58 \times 10^{-9}$ | | intronic | | | *CELF4* |
| rs12938775 | -0.0202 | 0.0035 | $8.54 \times 10^{-9}$ | SWB | intronic | | | *PAFAH1B1* |
| rs12961969 | 0.0253 | 0.0045 | $2.16 \times 10^{-8}$ | | intergenic | | | *CELF4* |
| rs35688236 | 0.0213 | 0.0038 | $2.35 \times 10^{-8}$ | | intronic | | | *PDCD6IP* |
| rs2150462 | -0.0217 | 0.0039 | $2.66 \times 10^{-8}$ | SWB | intergenic | | | *ELAVL2* |
| rs12903563 | 0.0198 | 0.0036 | $2.86 \times 10^{-8}$ | | intronic | | | *LINGO1* |

**Panel B. Additional DS and Neuroticism SNPs Identified Via Proxy-Phenotype Analysis** **SWB ( $N = 278,956$ ) → DS ( $N_{eff} = 111,106$ )**
| SNP | Beta<sub>DS</sub> | SE<sub>DS</sub> | $p$-value<sub>DS</sub> | Location | Non-syn | eQTL | Nearest protein-coding gene |
| --- | --- | --- | --- | --- | --- | --- | --- |
| rs4481363 | 0.0136 | 0.0038 | $3.06 \times 10^{-4}$ | intronic | | | *ZNF184* |
| rs6904596 | -0.023 | 0.0059 | $9.79 \times 10^{-5}$ | exonic | | Y | *MAT2B* |

**SWB ( $N = 239,982$ ) → Neuroticism ( $N = 131,867$ )**
| SNP | Beta<sub>Neur</sub> | SE<sub>Neur</sub> | $p$-value<sub>Neur</sub> | Location | Non-syn | eQTL | Nearest protein-coding gene |
| --- | --- | --- | --- | --- | --- | --- | --- |
| rs4481363 | 0.0151 | 0.0040 | $1.86 \times 10^{-4}$ | intronic | | | *ZNF184* |
| rs6904596 | -0.0264 | 0.0072 | $2.49 \times 10^{-4}$ | exonic | | Y | *MAT2B* |
| rs10838738 | 0.0178 | 0.0039 | $5.03 \times 10^{-6}$ | intronic | | Y | *MTCH2* |
| rs10774909 | -0.015 | 0.0039 | $1.20 \times 10^{-4}$ | intronic | | | *NOS1* |

We further investigated the biological mechanisms underlying the GWAS results by applying stratified LD Score regression (*27*) to our meta-analysis results. The method takes as given some categorization of SNPs, and for each category, it estimates the expected increase in the explanatory power of a SNP due to its being in that category. In our first analysis, we report estimates for all 53 functional categories included in the “baseline model” (tables S12.1-12.3); the results for SWB, DS, and neuroticism are broadly similar and are in line with what has been found for other phenotypes (*27*). In our second analysis, the categories are groupings of SNPs likely to regulate gene expression in cells of a specific tissue. The estimates for SWB, DS, and neuroticism are shown in fig. 3a. As a benchmark, we also provide results from the GIANT consortium’s most recent study of height (*28*). The height results are clearly distinguished by the much higher estimates for the categories connective/bone, cardiovascular, and skeletal muscle. In contrast, the estimates for the category central nervous system are significantly positive for all three of our phenotypes, and the estimates for adrenal/pancreas for two of the three, SWB and DS. This latter result is in line with previous research on the role of hypothalamic-pituitary-adrenal (HPA)-axis in depressed patients (*29*).

**Fig. 3.**
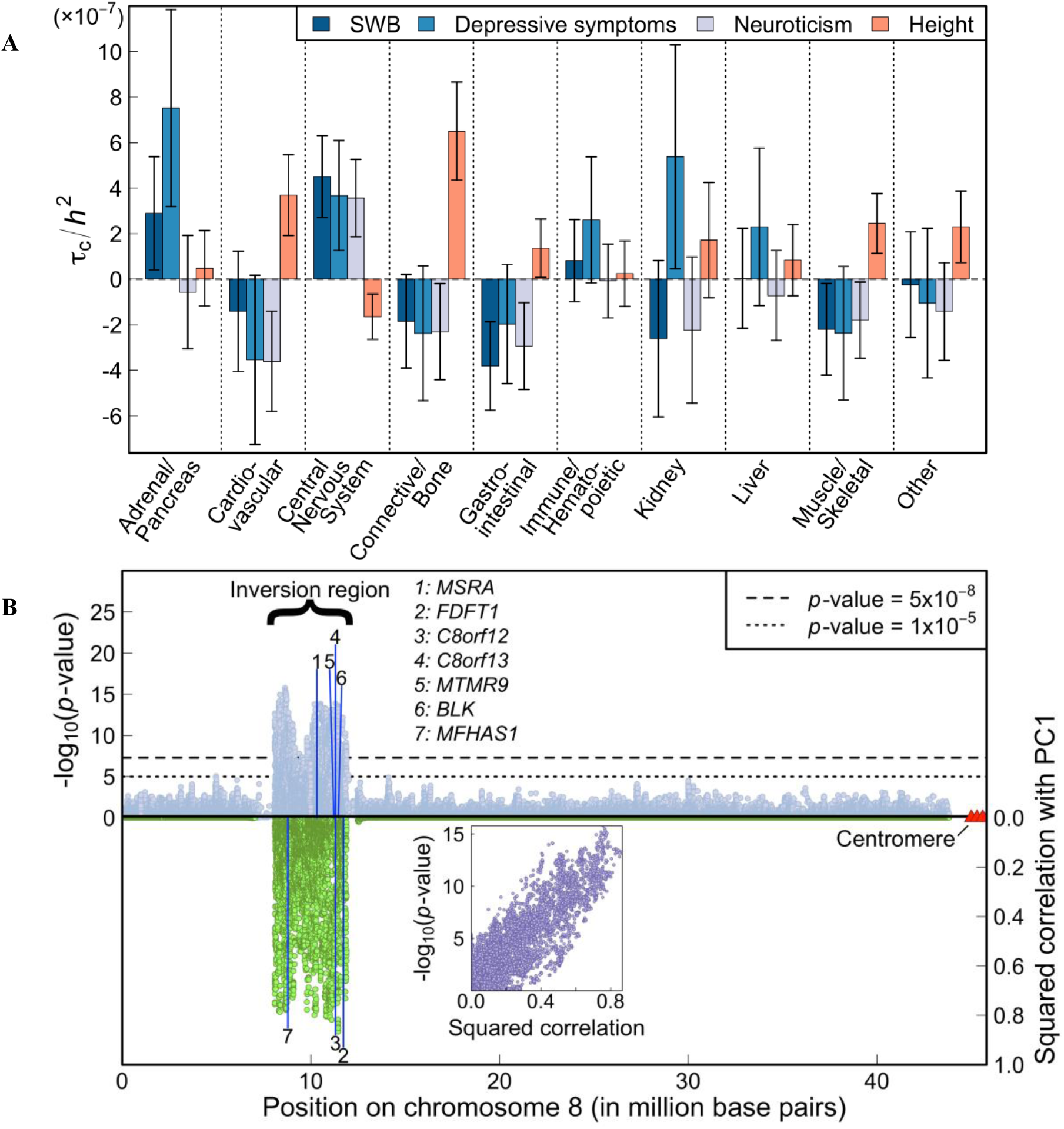
Results from selected biological analyses. **(A)** Estimates of the expected increase in the phenotypic variance accounted for by a SNP due to the SNP’s being in a given category (*τ_c_*), divided by the LD Score heritability of the phenotype (*h^2^*). Each estimate of *τ_c_* comes from a separate stratified LD Score regression, controlling for the 52 functional annotation categories in the “baseline model.” The bars represent 95% CIs (not adjusted for multiple testing). To benchmark the estimates, we compare them to those obtained from a recent study of height (*25*). **(B)** Inversion polymorphism on chromosome 8 and the 7 genes for which the inversion is a significant *cis*-eQTL at FDR < 0.05. The upper half of the figure shows the Manhattan plot for neuroticism for the inversion and surrounding regions. The bottom half shows the squared correlation between the SNPs and the principal component that captures the inversion. The inlay plots the relationship, for each SNP in the inversion region, between the SNP’s significance and its squared correlation with the principal component that captures the inversion.

We also investigated potential molecular mechanisms underlying the effects of the inversion on chromosome 8. The chromosome 8 inversion, and the 7 genes for which we find the inversion is a significant *cis*-eQTL at FDR < 0.05, are depicted in fig. 3b. The fact that these genes are all positioned in close proximity to the inversion breakpoints suggests that the molecular mechanism could involve the relocation of regulatory sequences. Three of the genes (*FDFT1*, *MSRA*, *MTMR9*) are known to be highly expressed in tissues and cell types that belong to the central nervous system, and two (*BLK, MFHAS1*) in the immune system.

To summarize, we report a number of credible genetic associations with our primary phenotype of SWB as well as our auxiliary phenotypes of DS and neuroticism. The small effect sizes help shed light on why earlier studies of these phenotypes have had limited success. For example, given our winner’s-curse-adjusted estimates of the effect sizes, we calculate that the power for detecting the genome-wide significant SNPs from our analysis in the largest previous GWAS of depression and neuroticism was only ~0.012% and ~0.00005%, respectively (table S11.2).

Our study found many more credible associations than prior work due to two strategies. First, our GWAS had much larger sample sizes. Our findings support the view that GWAS can succeed even for highly polygenic phenotypes once sample sizes are large enough (*9, 30*). Second, our proxy-phenotype strategy exploited the strong genetic overlap between SWB, depression, and neuroticism. We anticipate that future efforts to identify genetic variants associated with these phenotypes could similarly benefit from studying them in concert.

## Acknowledgements

This research was carried out under the auspices of the Social Science Genetic Association Consortium (SSGAC). The SSGAC seeks to facilitate studies that investigate the influence of genes on human behavior, well-being, and social-scientific outcomes using large genome-wide association study meta-analyses. The SSGAC also provides opportunities for replication and promotes the collection of accurately measured, harmonized phenotypes across cohorts. The SSGAC operates as a working group within the CHARGE consortium. This research has also been conducted using the UK Biobank Resource. The study was supported by funding from the U.S. National Science Foundation (EAGER: “Workshop for the Formation of a Social Science Genetic Association Consortium”), a supplemental grant from the National Institute of Health Office of Behavioral and Social Science Research, the Ragnar Söderberg Foundation (E9/11), the Swedish Research Council (421-2013-1061), The Jan Wallander and Tom Hedelius Foundation, an ERC Consolidator Grant (647648 EdGe), the Pershing Square Fund of the Foundations of Human Behavior, and the NIA/NIH through grants P01-AG005842, P01-AG005842-20S2, P30-AG012810, and T32-AG000186-23 to NBER and R01-AG042568-02 to the University of Southern California. We are grateful to Peter M. Visscher for advice, support, and feedback. We thank Samantha Cunningham and Nishanth Galla for research assistance. A full list of acknowledgments is provided in the supplementary materials. The authors declare no competing financial interests. Upon publication, results can be downloaded from the SSGAC website (http://thessgac.org/). Data for our analyses come from many studies and organizations, some of which are subject to a MTA, and are listed in the Supplementary Information.

